# Application of stable isotopes to study movement ecology and diet variation in a migratory songbird

**DOI:** 10.1101/2021.07.10.451926

**Authors:** Andrea Contina, Allison K. Pierce, Scott W. Yanco, Eli S. Bridge, Jeffrey F. Kelly, Michael B. Wunder

**Affiliations:** University of Colorado Denver, Department of Integrative Biology, Denver, CO, USA; Department of Integrative Biology, University of Texas at Austin, Austin, TX, USA; Center for Biodiversity and Global Change, Yale University, New Haven, CT, USA; Department of Ecology and Evolutionary Biology, Yale University, CT, USA; Oklahoma Biological Survey, University of Oklahoma, Norman, OK, USA; Corix Plains Institute and Department of Biology, University of Oklahoma, Norman, OK, USA; Department of Biology, University of Oklahoma, Norman, OK, USA

## Abstract

Whether and how migratory organisms exhibit inter-individual behavioral and/or physiological variation across movement strategies remains an open question. The Dark-eyed Junco (Junco hyemalis) is a migratory songbird known for its intra-species variation displayed in relation to morphology, song repertoires, and migration. Thus, studies focusing on juncos can reveal how migratory strategy may covary with other individual-scale factors and, therefore, identify the selective forces driving intra-species variations throughout its distribution. We used Dark-eyed Junco hydrogen stable isotope feather values (delta2H) and implemented a Bayesian framework to infer the breeding and molting origin of migratory juncos captured on their winter grounds in Oklahoma, United States (U.S.). We modeled the distribution of feather hydrogen stable isotope values as a function of five morphological variables including body mass and fat deposition measured in Oklahoma during the winter. We then investigated the trade-off between longer and more energetically costly migration strategies in relation to diet preferences through carbon (delta13C) and nitrogen (delta15N) stable isotope analysis from feather values. Dark-eyed Juncos wintering in south central U.S. likely originate from multiple breeding populations in northern U.S. and Canada. Body condition at the wintering ground (e.g., mass) had no effect on feather hydrogen stable isotope abundance. However, we found a positive correlation between nitrogen and hydrogen stable isotopes, suggesting that a trophic level shift towards insect consumption might occur in individuals migrating from southern latitudes. Increased insect-derived protein consumption might be explained by reduced fatty-acid reserves necessary to complete a shorter migratory journey.

## Introduction

Animal migration is a multifaceted life-history trait that evolved across taxa through complex behavioral and physiological adaptations (Alerstam and Bäckman, 2018). Migratory species maintain energetic balance between fuel consumption (e.g., fatty acid catabolism) and replenishing resources to effectively perform a long-distance migration (Guglielmo, 2018). However, research into how migrants sustain themselves during migration remains particularly challenging due to the small body size of many migratory songbirds and the fact that they travel, in some cases, thousands of kilometers per day (Rani et al., 2017; DeLuca et al., 2015). Species have developed unique behavioral strategies associated with seasonal movements (Horton et al., 2016), which depend on resource availability and seasonal environmental stochasticity. Thus, an understanding of these strategies requires examination of fluctuations in body mass and fat deposition in conjunction with behavior across the spatial extent of the species’ distribution. This is relevant because variation in migration and foraging behavior at the intra-species level can place different evolutionary pressures on individual physiology and act as selective forces across populations. That is, there might be a trade-off between longer and more energetically costly migratory routes, potentially resulting in decreased body condition, and alternative strategies involving diet changes or morphological adaptations (e.g., longer or pointed wings) to accommodate the increased energetic demands of their migratory journey (Kaboli et al, 2007; Reif et al., 2016).

Many species of songbirds are too small to be fitted with tracking devices (e.g., satellite transmitters) and information on their migratory patterns remain unknown. However, indirect molecular approaches can be implemented to study large-scale movements without the use of extrinsic tracking devices. (Rundel et al., 2013). In particular, movement ecology studies have exploited the latitudinal gradient in stable hydrogen isotopes occurring across North America to address questions related to avian migratory patterns and connectivity between breeding and wintering grounds (Rubenstein and Hobson, 2004; Vander Zanden et al. 2018; Besozzi et al. 2021). Stable isotope ratios of hydrogen obtained from inert animal tissues (e.g., claws, hair, and feathers), provide insight on the environmental conditions and latitudinal gradient where the keratin-based tissues were grown and offer a valid molecular tool to infer large-scale animal movements in species too small for direct movement observations (Hobson, 2005). Importantly, most songbirds replace their flight feathers at the breeding ground and before starting fall migration to minimize the energetic costs of the journey (Barta et al., 2008). By replacing old wing feathers with new ones, presumably in better conditions, migratory birds gain aerodynamic efficiency (Bowlin and Wikelski, 2008). Thus, feather samples collected at the wintering grounds, sometimes thousands of kilometers away from the breeding locations, can be used to infer where the feathers were grown based on stable isotope similarities between environmental and tissue hydrogen values (Wunder, 2010).

A suitable study system for disentangling different migratory strategies from other key evolutionary adaptations is offered by the Dark-eyed Junco (*Junco hyemalis*), a migratory songbird comprising numerous subspecies broadly distributed across North America. It is found breeding and molting in the northern regions of Canada and Alaska, and mountainous regions of the United States (U.S.) and wintering in the lower latitudes of Canada and the majority of the continental U.S. and northern Mexico (Nolan et al., 2002). This species has shown negative population trends of over 40% in some areas in the last decade (PIF 2017; Rockwell et al., 2019) and its physiology and migratory behaviors have been the focus of numerous studies (Nolan and Ketterson, 1983; Rogers et al., 1994.; Fudickar et al., 2016; Liebgold et al., 2019).

Bridge et al. (2010) analyzed hydrogen isotope ratios from secondary feathers in a population of Dark-eyed Juncos, all from the same *hyemalis* subspecies group (slate-colored), sampled during the winter of 2009 in central Oklahoma (U.S.) and showed isotopic variation ranging from -175.6‰ to -118.6‰ from Vienna Standard Mean Ocean Water (VSMOW). This large range of variation (∼60‰) in hydrogen stable isotope values prompted the authors to speculate that individuals from a wide geographic range of breeding, and consequently molting, locations wintered in Oklahoma, although the study did not include explicit probabilistic geographic assignments of wintering individuals, nor did it consider their variation in trophic level.

Although the subspecies of Dark-eyed Junco’s are primarily differentiated by morphology and/or geographic distribution, considerable behavioral diversity between and even within subspecies presents a unique opportunity to address large-scale spatial and ecological questions. We used hydrogen stable isotope data to estimate breeding location origins of a population of Dark-eyed Juncos wintering in Oklahoma, and we collected a suite of morphological measurements after the birds completed fall migration. Moreover, we expect that Oklahoma wintering birds originating from northern latitudes would have a higher carbohydrate intake, but lower protein consumption compared to wintering birds from southern latitudes due to their need to fuel a longer migratory journey. To test this hypothesis, we assessed pre-migratory diets by examining carbon and nitrogen stable isotope ratios in flight feathers. In addition to quantifying geographic connectivity in the Dark-eyed Junco population wintering in Oklahoma, this study offers new insights into how this species negotiates tradeoffs between migration distance and diet. Importantly, we present a framework to identify possible confounding effects in explanatory models for different migration strategies across populations (Yanco et al. 2020).

## Methods

### Study Area

We studied Dark-eyed Juncos wintering in central Oklahoma where the species forages in open woodland habitats and feeds on grass sprouts, *Taraxacum* sp. and various species of insects (Bridge et al. 2010). The weather conditions during our study (January and February 2009) as recorded by the Oklahoma Climatological Survey and the Oklahoma Mesonet (www.mesonet.org) at the Norman (NRMN) weather station indicated monthly average temperature ranging from 2.7 to 8.9 °C, monthly average precipitation ranging from 22.8 to 24.6 mm, and monthly average wind speed ranging from 15.6 to 18.1 kph.

### Sample collection and stable isotope analysis

We used published stable hydrogen isotope data from 80 Dark-eyed Juncos sampled in central Oklahoma, USA (35°10′ N, 97°26′ W) during winter 2009 (Bridge et al. 2010). After removing individuals with missing capture information, our data set comprised stable hydrogen isotope values of secondary feathers from 76 migrants that we used to infer geographic breeding and molting origin. We also present novel carbon and nitrogen stable isotope data obtained from a subset (N = 74) of those migrants sampled in Oklahoma that we used to study habitat use and trophic level. However, after filtering out individuals with missing morphometric measurements we had data from 62 migrants (39 adult males and 23 adult females) that we used for morphological models. All samples were collected between January and February of the same year. Feathers were prepared for analysis of stable isotope ratios of hydrogen (δ²H), carbon (δ^13^C), and nitrogen (δ^15^N) by following standard washing protocol in a 2:1 solution of chloroform-methanol to remove debris and oil contaminants (Paritte and Kelly 2009; Chew et al. 2019). Samples for hydrogen stable isotope ratios analysis were processed through a Thermo-Finnigan Delta V isotope ratio mass spectrometer interfaced with a high-temperature pyrolysis elemental analyzer (TC/EA, Thermo-Finnigan, Bremen, Germany). Feather δ²H values (δ²H_f_) are reported as mean ± SD in delta notation of parts per mil (‰) from the standards (δD_sample_ = [(*R*_sample_/*R*_standard_) − 1]) compared to the Vienna Standard Mean Ocean Water (VSMOW). We used chicken feather standard (CFS; -147.4‰), cow hooves (CHS; -187‰), and bowhead whale baleen (BWB; -108‰) as keratin which are widely adopted for sample comparative equilibration (Wassenaar and Hobson, 2003). The details of hydrogen sample measurement are in Kelly et al. 2009. We conducted carbon and nitrogen stable isotope ratios analysis on a Thermo-Finnigan DeltaPlus isotope ratio mass spectrometer connected to a Carlo Erba elemental analyzer. We normalized the δ^13^C and δ^15^N values to the Vienna Pee Dee Belemnite (VPDB) and AIR scales, respectively, and used Brown-headed cowbird (BHCO; *Molothrus ater*) feather standards. The values of δ^13^C and δ^15^N standards were -15.7 ± 0.1‰ (δ^13^C BHCO), and 7.6 ± 0.1‰ (δ^15^N BHCO).

### Feather Calibration and Geographical Assignments

We implemented a Bayesian analytical framework in the R package assignR to spatially reconstruct juncos’ breeding and/or molting areas (Ma et al., 2020; R Core Team, 2020). First, we rescaled stable hydrogen isotope values in precipitation (δ²H_p_) from a North America geostatistical model (Bowen et al. 2005) to 22 known-origin Song Sparrow (*Melospiza melodia*) samples provided by Hobson et al. (2012) and Hobson and Koehler (2015). To ensure a more accurate isotopic calibration (Wunder 2010), we used linear regression to model mean tissue isotope values as a function of δ²H_p_ using the *calRaster* function in assignR. We recognize that a species-specific calibration provides higher degree of accuracy. However, when species-specific known-origin tissues are not available, calibration data from closely related species can be used (Hobson et al., 2012). Our data set comprised entirely unknown-origin feather samples from Dark-eyed Juncos; therefore, we used published data of *M. melodia*, a species of comparable size, diet, and spatial distribution (Arcese et al., 2002; Hobson et al., 2012).

We determined the molting locations of juncos wintering in Oklahoma by computing posterior probability density maps through the *pdRaster* function in assignR (Wunder 2010; Ma et al., 2020). We present normalized posterior probability maps representing both the individual and cumulative group probabilities of origin from each grid cell restricted to the breeding and year-round distribution ranges (BirdLife International 2016). To visualize the cumulative group probabilities of origin, we clustered the samples into four isotopic groups based on the reported stable isotope feather values (δ²H_f_) across the junco data set arranged from the lowest values (G1) to the highest values (G4): the first group (G1) included δ²H_f_ values ranging from -175.6‰ to -160.2‰ (M = -167.1‰, SD = 5.02‰, N = 18); the second group (G2) included values ranging from -159.3‰ to -150.4‰ (M = -154.9‰, SD = 3.17‰, N = 23); the third group (G3) included values ranging from -149.8‰ to -140.1‰ (M = -145.4‰, SD = 2.88‰, N = 19); and the fourth group (G4) included values ranging from -139.3‰ to -118.6‰ (M = -130.4‰, SD = 6.39‰, N = 16). Even though this clustering approach is *ad hoc*, it still represents a useful subdivision based on δ²H_f_ variation across birds in our dataset. Wassenaar and Hobson (2006) estimated that ±3‰ variation in hydrogen stable isotope extracted from feathers translates approximately to 1 degree of latitude (or ∼111 km). Therefore, the differential range of about 10‰ or more that we used to cluster birds into four groups corresponds to ∼370 km of geographic separation (Hobson and Wassenaar, 1996; Wassenaar and Hobson, 2006). This distance is presumably large enough to consider our samples from different breeding populations.

### Modeling morphometric parameters

We investigated the effect and relative predictive importance of five different morphometrics on mean hydrogen stable isotope ratios in Dark-eyed Junco feathers. Our explanatory variables were: wing chord (mm), tarsus length (mm), tail length (mm), mass (g), and fat depositions scored on a six-point ordinal scale (Helms and Drury 1960). We did not include body mass index normalized by a structural measure because this would be collinear with other included variables and additive model combinations should account for size normalized mass (Green, 2001). We controlled for morphological variance across males and females by including ‘sex’ as a random effect in all our models and to standardize evaluation of predictive influence between the morphometric variables measured on different scales we Z-scored values for all variables. We modeled a complete combinatorial set of linear models in the R package MuMIn (Barton and Barton, 2019) and computed the relative importance of individual morphometric variables through a variable importance analysis using Akaike Information Criterion corrected for small sample sizes (AICc) and summed model weights for each model containing a given variable. We then compared the sum of all the AICc weights and odds ratios to evaluate the relative importance of each variable.

### Trophic Level Analysis

We analyzed stable carbon (δ^13^C) and nitrogen (δ^15^N) isotopes in feather samples to explore trophic level partition among birds migrating from northern and southern latitudes as revealed by hydrogen (δ^2^H) data. We exploited the occurrence of distinct plant photosynthetic pathways, knowing that trees and shrubs in temperate climates (e.g., C3 plants) are depleted in carbon compared to plants inhabiting arid environments and adopting C4 or CAM photosynthesis (Keeley and Rundel, 2003). These differences help to assess the proportion of proteins deriving from C3 and C4 plants, while higher nitrogen values indicate a shift towards a diet less dependent on seeds and richer in insects. Thus, we used the *corr*.*test* function in the R package psych (Revelle and Revelle, 2015) to perform a Pearson correlation test between carbon (δ^13^C) and hydrogen (δ^2^H) and between nitrogen (δ^15^N) and hydrogen (δ^2^H) stable isotopes. This test highlights diet trends in relation to latitudinal origin as determined by δ^2^H data. We then used the *levelplot* function in the R package lattice (Sarkar 2008) to create a visual representation of δ^2^H values from feathers collected in Oklahoma and their distribution within the sampling space.

## Results

### Rescaling model and posterior probability density maps

The rescaling model provided a robust relationship between precipitation isotope values (δ²H_p_) and known-origin samples of the species *M. melodia* (y = -35.08 + 0.91x; R² = 0.79, CI_95%_ [0.74, 0.95], n = 22) and allowed us to calibrate our isoscape to determine junco’s posterior probability of origin (Fig 1 – panel A). We obtained a rescaled environmental isoscape raster of 52,884 cells at 0.33km resolution and a range of hydrogen stable isotope values spanning - 193.3‰ and -41.3‰ (M = -122.9‰; SD = 37.2‰; Fig 1 – panel B and C).

**Figure 1.**
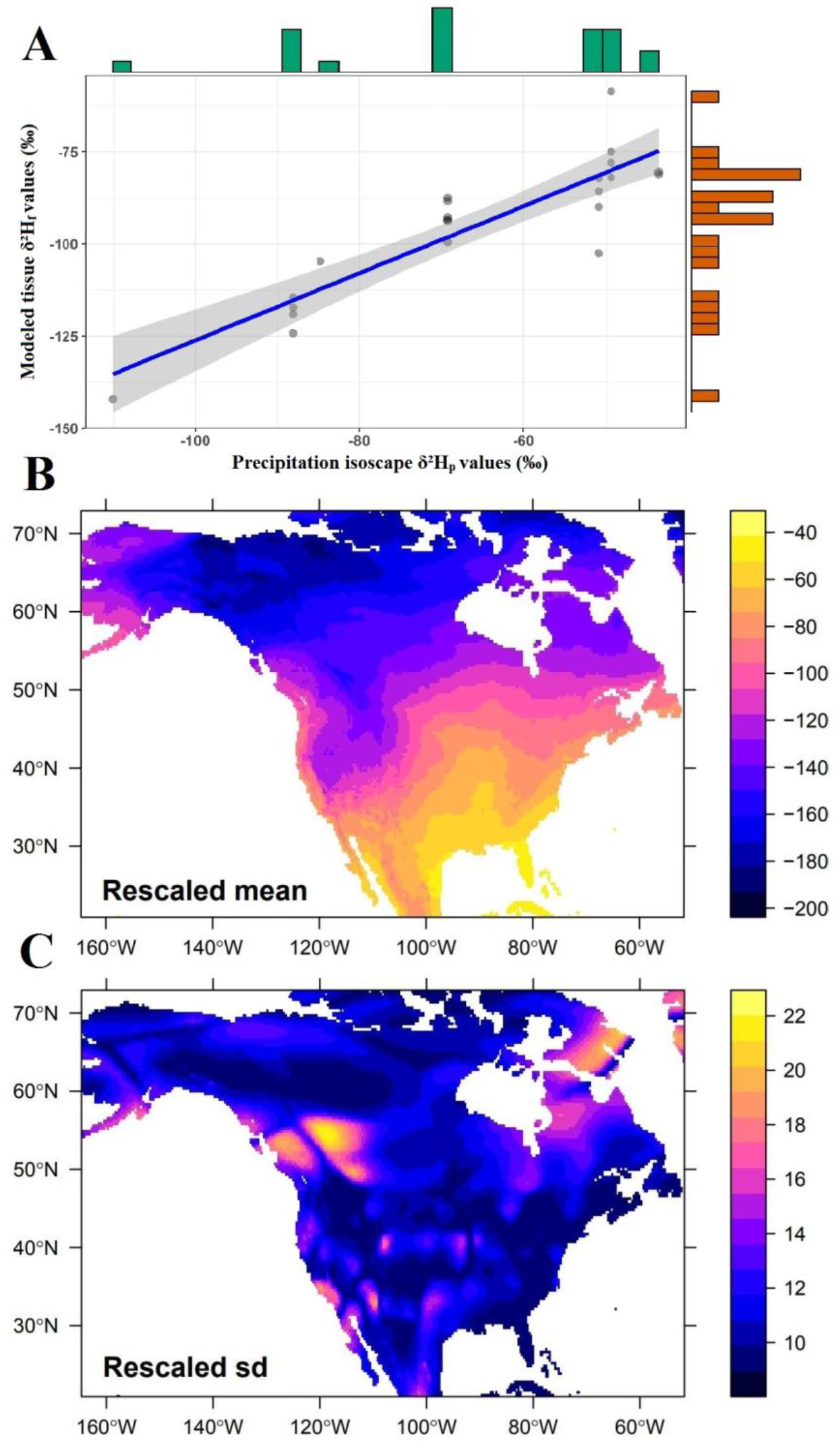
Linear regression model between environmental precipitation isotope values (δ²H_p_) and 22 known-origin feather samples of *M. melodia* samples (panel A) used to rescale the North America hydrogen isoscape (panel B and C) for assigning migratory juncos wintering in Oklahoma to their molting locations at the breeding grounds across northern United States and Canada. The green and orange bars along the horizontal and vertical axis represent the distributions of the sampling data across environmental and tissue isotope values, respectively (panel A). The calibrated hydrogen isoscape mean values and standard deviations are presented in panel B and C, respectively.

Once we performed isoscape tissue calibration, we inferred the spatial extent of the breeding and molting origin of 76 Dark-eyed Juncos wintering in Oklahoma through a Bayesian probabilistic framework. Our posterior probability density maps, show distinct latitudinal variation across the breeding and molting range of migratory juncos (Fig 2). Overall, the northernmost migrants were assigned to either Alaska and western Canada, including Yukon Territory and British Columbia, or central Northwestern Territories, while southern migrants were assigned to either Northern Saskatchewan or southern Manitoba and northwestern United States near the Rocky Mountains. However, even though the probability assignment surfaces for the southern migrants (group G3 and G4) were largely distributed at latitudes lower than 60°N (Fig 2), we also found weak probabilities of origin along the western coast of Alaska due to the underlying structure of the precipitation isoscape used for rescaling (Fig 1, panel B-C). Taken together, these results suggest that different Dark-eyed Junco populations admix into the same wintering area in central Oklahoma.

**Figure 2.**
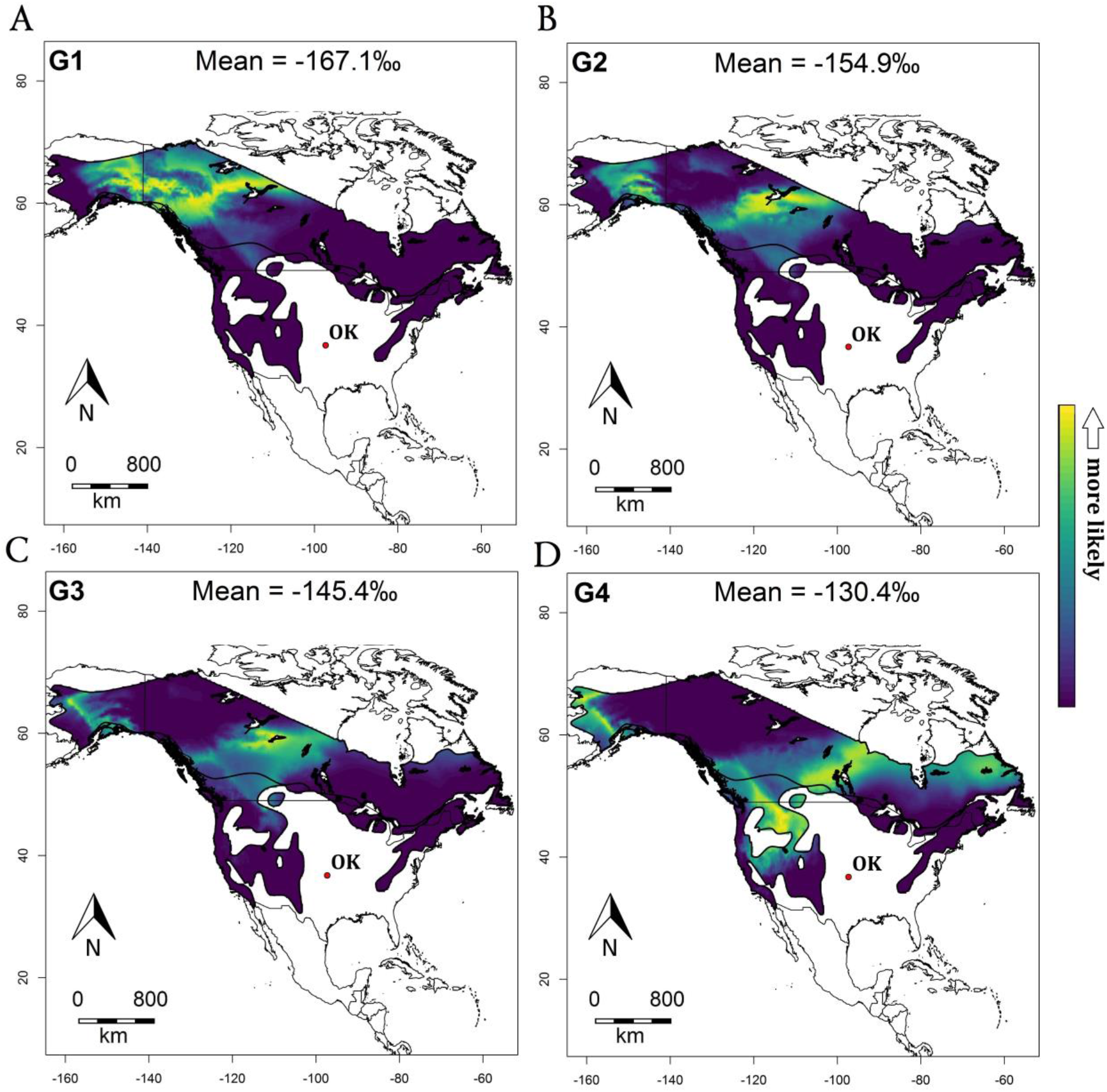
Cumulative assignment posterior probability density maps across four isotopic groups of 76 Dark-eyed Juncos wintering in Oklahoma (red dot) and breeding at northern latitudes in the United States and Canada. The bright yellow color represents high probabilities of molting origin. The northern migrants (groups G1 and G2; panel A and B) were assigned to either Alaska and western Canada, including Yukon and British Columbia, or central Northwestern Territories. The southern migrants (groups G3 and G4; panel C and D) were prevalently assigned to either Northern Saskatchewan or southern Manitoba and northern western United States. The hydrogen stable isotope approach suggests that Dark-eyed Junco migrants from different breeding populations converge into the same location in Oklahoma over the winter.

### Variable importance analysis

The analysis of the importance of variables revealed no clear predictive support for any variable considered in our models. We fit a full set of 32 linear mixed-effect models for mean δ²H_f_ including single variable models and all the possible combinations of the five predictors, with a random effect of sex and ranked them using Akaike Information Criterion corrected for small sample size (AICc). All 5 variables we considered had relatively high summed AICc evidence weights suggesting models containing each respective variable outperformed those that did not (Table 1 and Table 2). Fat score had the highest summed AICc weight value and odds ratios, indicating that models containing the fat score variable accounted for all of the weight of AICc values for the model set (Table 1). However, the estimated effect of fat score on mean δ²H_f_ was indistinguishable from 0 in all models weighted by AICc (Fig 3A) suggesting that despite being the most comparatively informative variable, it had no discernible effect on δ²H_f_ values in our data set. Similarly, estimates for other variables we considered also lacked support for non-zero effects Fig 3B).

**Table 1.** Variable importance results. Although the variable ‘body fat’ emerged as the comparatively best predictor of hydrogen variation in feathers, the effect for this and all other variables could not be differentiated from zero.

| Variable | Sum of AICc<br>model weights | Odds<br>ratio |
| --- | --- | --- |
| Fat | 1 | inf |
| Mass | 0.87 | 6.69 |
| Wing | 0.86 | 6.14 |
| Tail | 0.72 | 2.57 |
| Tarsus | 0.54 | 1.17 |

**Table 2.** Akaike Information Criterion corrected for small sample size (AICc) model selection results. The most parsimonious model was the full model including all five morphological variables but was equally competitive with a model omitting tarsus based on <2 unit difference in ΔAICc values. Simpler single term or intercept only models were generally outperformed by more complex models.

| Model ID | Model Variables | logLik | AICc | delta | weight |
| --- | --- | --- | --- | --- | --- |
| 32 | h ~ Fat + Mass + Tail + Tarsus + Wing + (1\|sex) | -222.1 | 471.48 | 0 | 0.3 |
| 24 | h ~ Fat + Mass + Tail + Wing + (1\|sex) | -223.74 | 471.8 | 0.33 | 0.25 |
| 28 | h ~ Fat + Mass + Tarsus + Wing + (1\|sex) | -224.61 | 473.54 | 2.06 | 0.11 |
| 20 | h ~ Fat + Mass + Wing + (1\|sex) | -226.25 | 473.95 | 2.47 | 0.09 |
| 30 | h ~ Fat + Tail + Tarsus + Wing + (1\|sex) | -225.43 | 475.16 | 3.69 | 0.05 |
| 22 | h ~ Fat + Tail + Wing + (1\|sex) | -226.94 | 475.33 | 3.86 | 0.04 |
| 16 | h ~ Fat + Mass + Tail + Tarsus + (1\|sex) | -225.65 | 475.61 | 4.13 | 0.04 |
| 8 | h ~ Fat + Mass + Tail + (1\|sex) | -227.18 | 475.83 | 4.35 | 0.03 |
| 12 | h ~ Fat + Mass + Tarsus + (1\|sex) | -227.46 | 476.38 | 4.91 | 0.03 |
| 4 | h ~ Fat + Mass + (1\|sex) | -228.98 | 476.68 | 5.21 | 0.02 |
| 26 | h ~ Fat + Tarsus + Wing + (1\|sex) | -227.69 | 476.85 | 5.37 | 0.02 |
| 18 | h ~ Fat + Wing + (1\|sex) | -229.2 | 477.11 | 5.63 | 0.02 |
| 14 | h ~ Fat + Tail + Tarsus + (1\|sex) | -229.25 | 479.96 | 8.48 | 0 |
| 6 | h ~ Fat + Tail + (1\|sex) | -230.84 | 480.39 | 8.91 | 0 |
| 10 | h ~ Fat + Tarsus + (1\|sex) | -232.01 | 482.74 | 11.26 | 0 |
| 31 | h ~ Mass + Tail + Tarsus + Wing + (1\|sex) | -233.91 | 483.9 | 12.42 | 0 |
| 2 | h ~ Fat + (1\|sex) | -233.92 | 483.91 | 12.44 | 0 |
| 23 | h ~ Mass + Tail + Wing + (1\|sex) | -235.4 | 484.32 | 12.85 | 0 |
| 27 | h ~ Mass + Tarsus + Wing + (1\|sex) | -236.3 | 486.13 | 14.66 | 0 |
| 19 | h ~ Mass + Wing + (1\|sex) | -237.79 | 486.64 | 15.16 | 0 |
| 15 | h ~ Mass + Tail + Tarsus + (1\|sex) | -237.14 | 487.81 | 16.33 | 0 |
| 29 | h ~ Tail + Tarsus + Wing + (1\|sex) | -237.26 | 488.04 | 16.56 | 0 |
| 7 | h ~ Mass + Tail + (1\|sex) | -238.62 | 488.32 | 16.84 | 0 |
| 21 | h ~ Tail + Wing + (1\|sex) | -238.76 | 488.6 | 17.12 | 0 |
| 11 | h ~ Mass + Tarsus + (1\|sex) | -238.93 | 488.93 | 17.45 | 0 |
| 3 | h ~ Mass + (1\|sex) | -240.4 | 489.51 | 18.03 | 0 |
| 25 | h ~ Tarsus + Wing + (1\|sex) | -239.41 | 489.89 | 18.41 | 0 |
| 17 | h ~ Wing + (1\|sex) | -240.9 | 490.51 | 19.03 | 0 |
| 13 | h ~ Tail + Tarsus + (1\|sex) | -240.75 | 492.58 | 21.1 | 0 |
| 5 | h ~ Tail + (1\|sex) | -242.43 | 493.56 | 22.09 | 0 |
| 9 | h ~ Tarsus + (1\|sex) | -243.45 | 495.6 | 24.13 | 0 |
| 1 | h ~ (1\|sex) | -245.58 | 497.58 | 26.1 | 0 |

**Figure 3.**
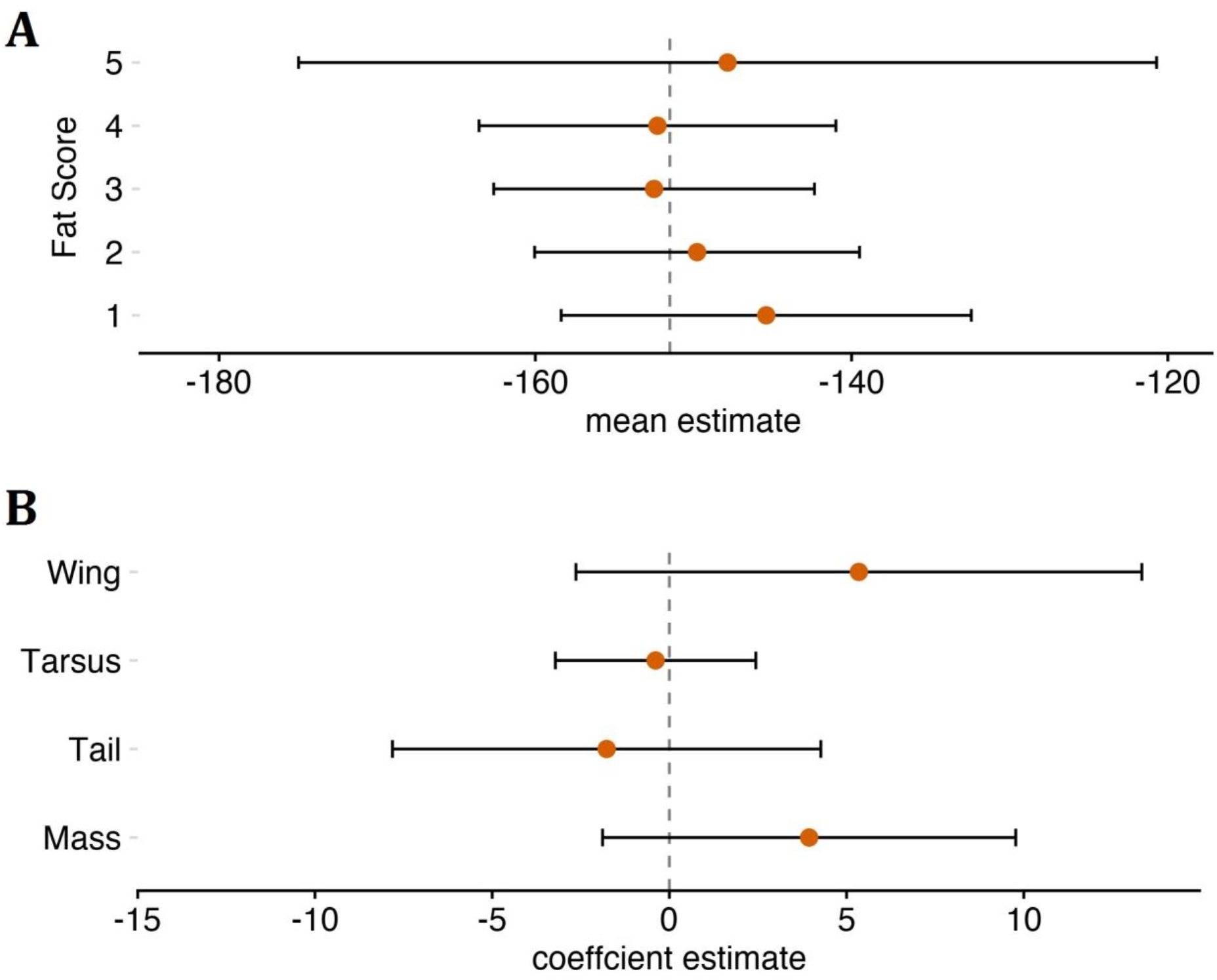
Model averaged estimates of mean δ²H_f_ values with 95% CI for levels of fat scores with centered and scaled morphometric metric measures held at 0, their mean value (A) and model averaged coefficient estimates of centered and scaled morphometric parameters with 95% CI with level 1 of fat score as the reference for the intercept (B). Both plots indicate effects for fat scores and morphometric variables on mean δ²H_f_ are insignificant for our data set as evidenced by CI overlap between fat score levels (A) and with zero (B).

### Migrant Diet Preferences and Trophic Level

The Pearson test did not identify a strong correlation between carbon (δ^13^C) and hydrogen (δ^2^H) isotopes values (r = -0.17, p = 0.22), but it suggested a moderately positive correlation between nitrogen (δ^15^N) and hydrogen (δ^2^H) isotopes values (r = 0.33, p = 0.01). These results suggest that a trophic level shift towards insect consumption might occur in migrants originating from lower latitudes (Fig 4 panel A and B). Interestingly, when samples were grouped by wing length (Fig 4 panel C and D), it appeared that birds with longer wings had a broader range of δ^13^C values, indicating variability in their habitat use. Moreover, when plotting δ^13^C against δ^15^N over δ^2^H feather isoscape within the sampling space, the changes in dietary niche across latitudinal origin appeared more evident (Fig 5 panel A-C; supplementary material; S1-S2). Northern migrants showed lower protein consumption (δ^15^N: 1-4‰) and C3-shrub habitat use (δ^13^C: -22 ? -24‰), while southern migrants were characterized by a higher trophic level (δ^15^N: 4-8‰) and δ^13^C towards C3-vegetation compatible with temperate forests (δ^13^C: -24 ? -26‰). Analysis of diet preferences between sexes did not show marked differences, although males seem to have a slightly larger dietary niche compared to females.

**Figure 4.**
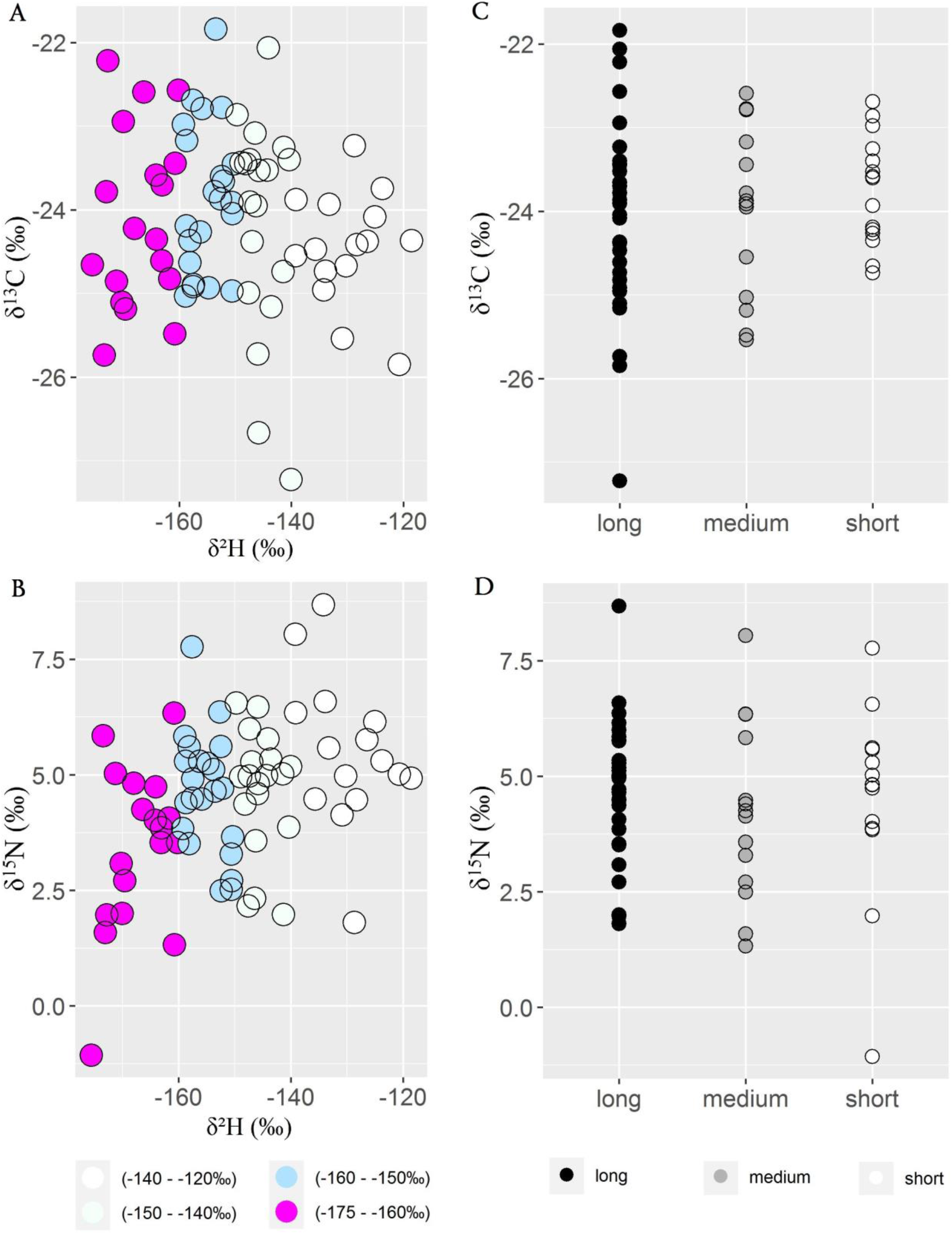
Correlation between δ^13^C and hydrogen δ^2^H (panel A) and nitrogen δ^15^N and hydrogen δ^2^H values (panel B). Migrants exhibiting longer wings show a broader range of δ^13^C values compared to medium-and short-length wings birds suggesting more flexibility in their habitat use (panel C). We did not find noticeable trends between δ^15^N and wing length (panel D).

**Figure 5.**
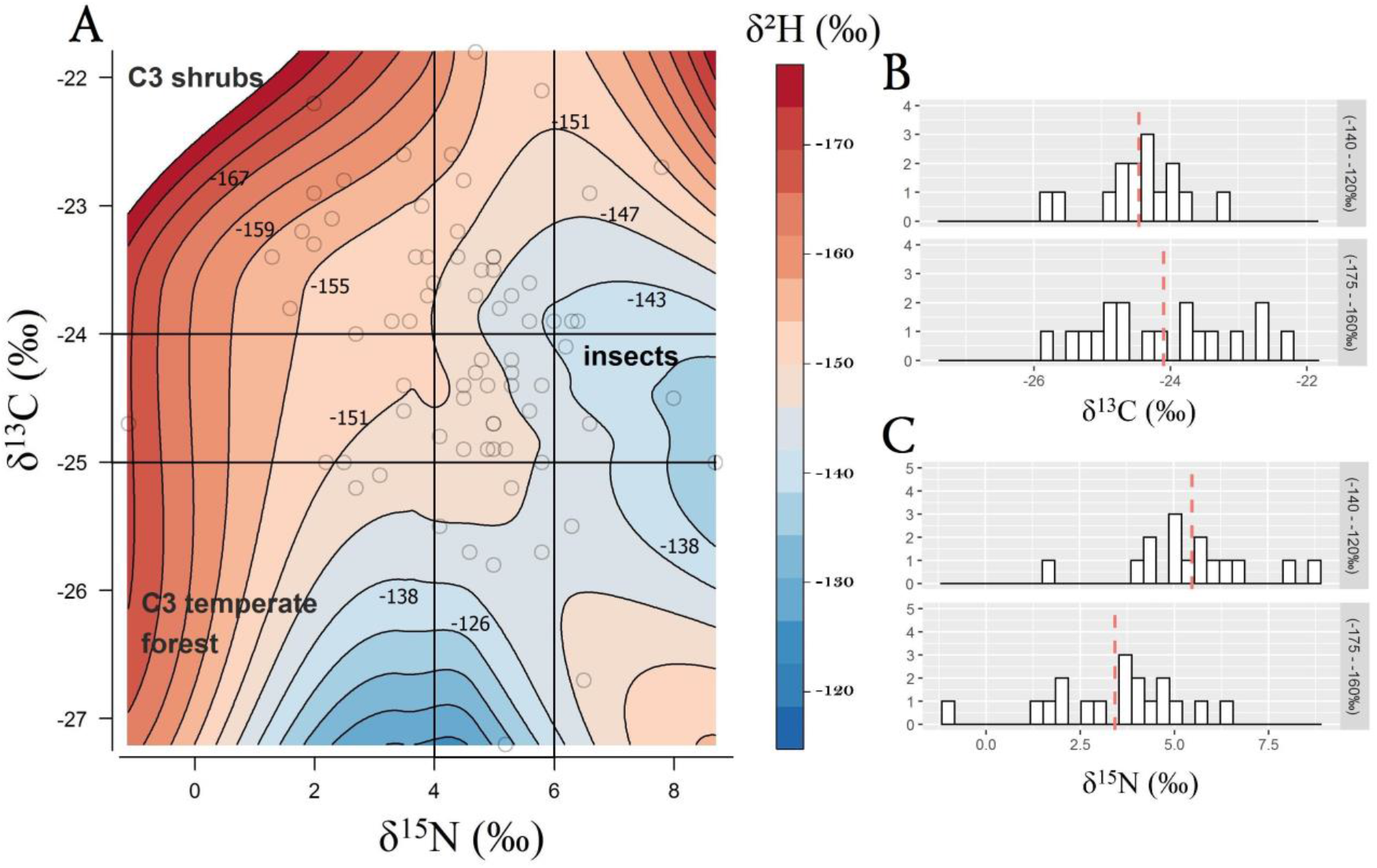
Isotopic diet composition of δ^13^C and δ^15^N plotted over δ^2^H spatial data distribution (panel A). Blue areas represent lower latitudes of origin (δ^2^H = from -110‰ to -145‰) while red areas represent higher latitudes of origin (δ^2^H = from -145‰ to -180‰). Each open circle represents a single individual. Northern migrants show a moderate trend towards lower protein consumption (δ^15^N: 1-4‰) and C3-plant habitat use (δ^13^C: -22 ? -24‰). Southern migrants show higher trophic level (δ^15^N: 4-8‰) and temperate forests habitat use (δ^13^C: -24 ? -26‰). Panels B and C show the distributions of δ^13^C and δ^15^N values across migrants originating from extreme northern and extreme southern latitudes. Dashed red lines represent isotope mean values for each group.

## Discussion

Bridge et al. (2010) conducted a captive experiment in which wintering Dark-eyed Juncos were fed *ad libitum* to demonstrate that fat depositions allow them to initiate migration earlier in the spring compared to the control group. We extended previous work by providing a geostatistical approach illustrating the explicit spatial distribution of breeding and molting origin of fall migrants and related that distribution to morphological characteristics and trophic level differentiation. Our geographical assignments clearly suggested that migratory juncos wintering in central Oklahoma originated from different breeding and molting areas (Fig 2). We identified northern and southern migrants while each group also showed intra-cohort spatial variation ranging from Alaska and western Canada to central Northwestern Territories, and Northern Saskatchewan to southern Manitoba and northwestern United States, respectively. However, despite the posterior probability surfaces showing population admixture at the wintering area in Oklahoma, our modeling effort showed no association between winter body condition and breeding or molting latitude.

Stable isotope ratios of hydrogen, carbon, and nitrogen, extracted from inert animal tissues, such as claws and feathers, reflect the isotope ratios in the environment where those tissues were grown and can provide inferences about large-scale animal movements and diet preferences in species with small body size (Rubenstein and Hobson, 2004; Zenzal et al., 2018; Vander Zanden et al., 2018). However, a clear advantage of adopting a multi-element stable isotope approach is that migrants can be assigned to different breeding and molting grounds of origin while their diet preferences are also revealed and clustered in trophic guilds (Herrera et al., 2003). The positive correlation between nitrogen (δ^15^N) and hydrogen (δ^2^H) values, although moderate, corroborated our hypothesis that wintering birds originating from northern latitudes would have lower protein intake relative to southern migrants and prefer a diet rich in carbohydrates before migrating. That is, increased carbohydrate anabolism and lower protein consumption is necessary to quickly synthesize enough subcutaneous lipid reserves to sustain a long-distance migration (Cecere, et al., 2011; Domer et al., 2018). Alternatively, higher δ^15^N values in southern migrants might be explained by environmental stressors. An early study in captive quails (*Coturnix japonica*) and geese (*Anser rossii*) showed that increased δ^15^N values can be obtained from growing tissues of individuals maintained under severe nutritional stress conditions (e.g., fasting) compared to control groups (Hobson et al. 1993). However, subsequent analysis in sparrows (*Melospiza melodia*) found no differences in δ^15^N between birds kept under a restricted diet versus birds fed *ad libitum* (Kempster et al., 2007) suggesting a less straightforward relationship between physiological stress conditions and δ^15^N values. Therefore, even though we cannot pinpoint the exact causes of increased δ^15^N values in growing feathers (e.g., baseline difference) and that other species have shown variation in feather δ^15^N values between boreal and more southern food webs based on land use (Hobson 1999), it is also possible that migratory juncos originating from southern latitudes were foraging at a higher level in the trophic system and consuming more insects compared to northern migrants. We do not exclude that such diet preference might be linked to an opportunistic foraging strategy due to an outburst of insect availability (Yang et al., 2008). However, analysis of isotopic niche shifts in avian migratory populations have shown sharp changes during different life cycle stages (Hahn, et al., 2013), confirming that variations in diet compositions have a defined role in maintaining fluctuating energy needs (Marshall et al., 2016). Moreover, our δ^13^C results indicated that northern juncos might exploit food resources from a broader range of habitats compared to southern juncos, ranging from pine forest to deciduous forest (e.g., oak) and C3 shrubs. Bearhop et al. (2004) studied the Thick-billed Vireos (*Vireo crassirostris*) in the Bahamas and based on mean claw δ^13^C values found differences among two groups of birds occupying environments with either prevalent xerophytic shrubs or pine forest. While our feather δ^13^C results showed that northern juncos seed consumption fell within the spectrum of plants characterized by a C3 photosynthetic pathway, and we found enough variation to highlight habitat occupancy differentiations across latitudinal origins (Fig 4 and Fig 5), we note that a robust environmental isotopic baseline is necessary to determine dietary and habitat preferences with confidence (Post, 2002).

The interactions between avian body mass fluctuations and migratory behaviors within and between seasons remain poorly understood (Brown and Sherry, 2006; Tsvey et al., 2007; Marra et al. 2015). Importantly, body condition appears to both influence and be influenced by migratory behavior. For example, body condition may affect migratory decisions (Schaub and Jenni 2000); several species of migrating shorebirds underwent longer migratory flights when departing with higher body condition scores (Anderson et al. 2019). Similarly, spring arrival timing of red knots (*Calidris canutus*) was associated with better body condition at a major stopover site (Duijns et al. 2017) and conditions experienced during winter months by American Redstarts (*Setophaga ruticilla*) lead to variation in body conditions that predicted spring departure dates (Marra et al., 1998). In our investigation, individuals that migrated minimum distances did non exhibit larger body masses or higher fat scores over the early winter period. However, our study was limited because individual migrants were not captured immediately upon arrival in Oklahoma. While challenging, an earlier capturing effort might reveal mass as a predictor of latitude of origin because the body condition of those migrants would provide a stronger indication of the migration strategy performed. Moreover, there may be other mechanisms that could explain the absence of a relationship between body conditions and migration strategies in our study population. For example, the taxonomy of the genus *Junco* has been challenging for decades and is far from being resolved (Miller 1941; Nolan et al., 2002; Aleixandre et al., 2013). The species *Junco hyemalis* has been recently proposed to be phylogenetically reassessed based on extensive genomic data to determine if it needs to be split into four new species that also account for distinct phenotypic variation (Friis et al., 2016). Thus, in a complex migratory system, it is possible that in some populations different morphs of migrants characterized by a larger and smaller than average body size could come into contact at the winter ground and therefore confounding the effects of energy consumption during migration. Alternatively, long distance migrants could also improve their body conditions (e.g., increase mass at stopover sites) before arriving at their final wintering destination and reduce the difference in body mass compared to short distance migrants. However, if the stopover site is only used to refuel without growing new feathers, the stable isotope analysis will still detect the northern origin (e.g., molting ground) and correctly identify the migration latitudinal origin without revealing any geographical information on the stopover location.

## Conclusions

By implementing a Bayesian assignment framework, we offered a probabilistic geographic explanation of the junco’s feather isotopic variation across the species’ breeding and molting areas in the northern hemisphere and attempted to explain latitudinal origin in relation to morphological traits (Wunder, 2010; Alerstam, 2011; Maggini et al., 2016). In our system, none of the variables considered had a clear effect on feather hydrogen stable isotope values (used as a proxy for latitudinal origin) although we found indication of a possible higher trophic level shift in individuals migrating from southern latitudes. These findings promote the concurrent use of different stable isotope markers as a means of studying interconnected behavioral traits. Our results also highlight the need to carefully examine confounding effects in explanatory models in relation to the species’ natural history and possible differential migratory behaviors across populations.

## Abbreviations

NRMN: Norman; AICc: Akaike Information Criterion corrected for small sample sizes; G1: Group 1; G2: Group 2; G3: Group 3; G4: Group 4.

## Authors’ contributions

AC conceptualized the project, analyzed the data, and wrote the manuscript. AKP analyzed the data, reviewed, and edited the manuscript. SWY provided key feedback on data analysis, reviewed, and edited the manuscript. ESB collected the data and reviewed the manuscript. JFK collected the data and reviewed the manuscript. MBW supervised data analysis and reviewed the manuscript.

## Supplementary Material

**Figure S1.**
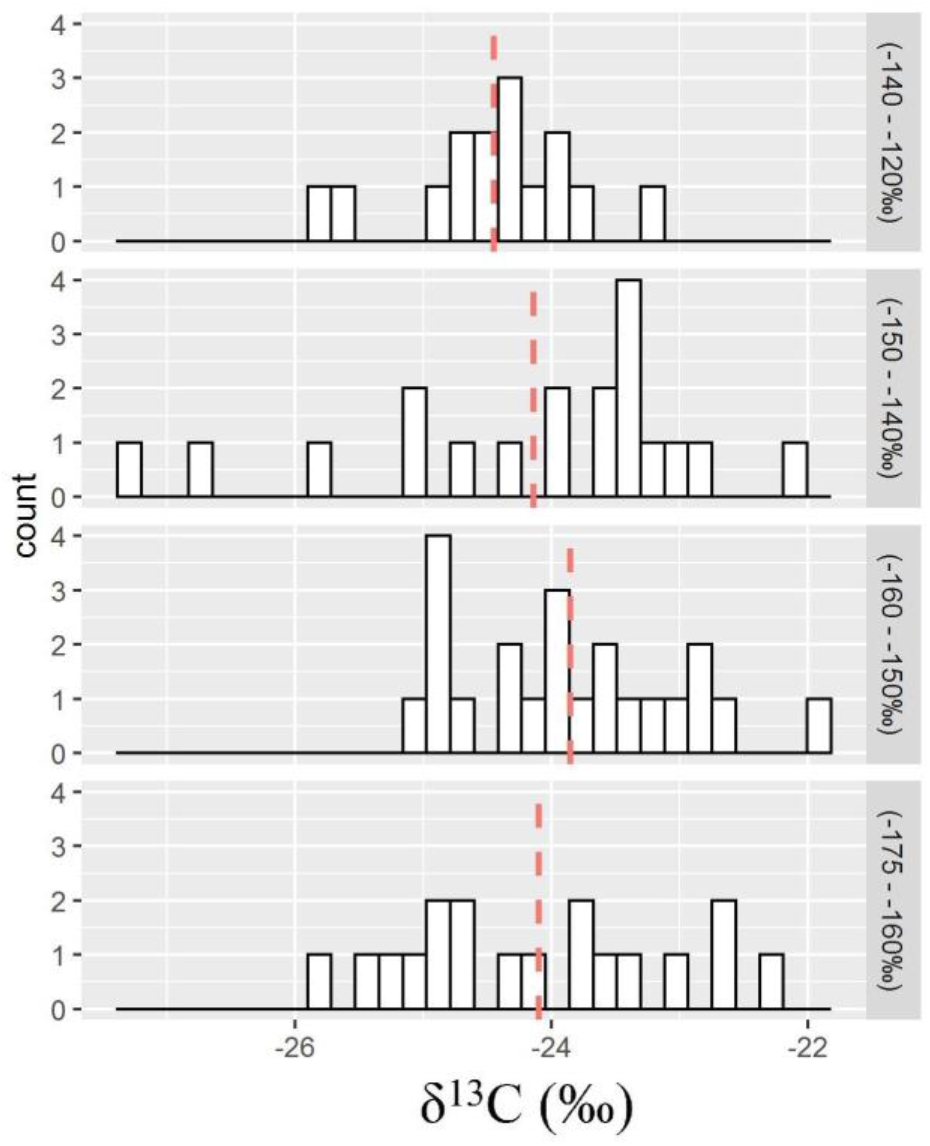
Distributions of δ^13^C values across migrants originating across latitudinal groups. Dashed red lines represent stable isotope mean values for each group.

**Figure S2.**
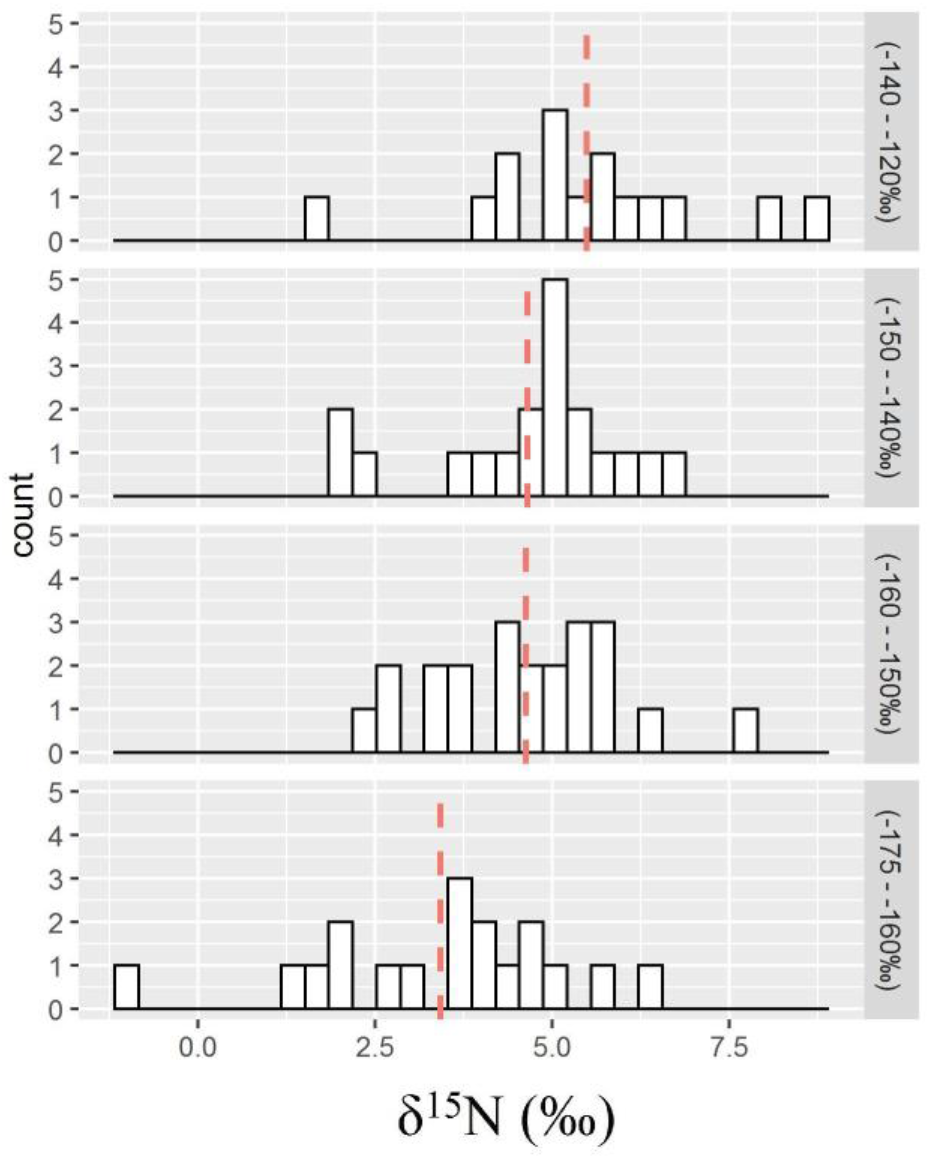
Distributions of δ^15^N values across migrants originating across latitudinal groups. Dashed red lines represent stable isotope mean values for each group.

